# Crowdfunded whole-genome sequencing of the celebrity cat Lil BUB identifies causal mutations for her osteopetrosis and polydactyly

**DOI:** 10.1101/556761

**Authors:** Mike Bridavsky, Heiner Kuhl, Arthur Woodruff, Uwe Kornak, Bernd Timmermann, Norbert Mages, 99 Lives Consortium, Darío G. Lupiáñez, Orsolya Symmons, Daniel M. Ibrahim

## Abstract

Rare diseases and their underlying molecular causes are often poorly studied, posing challenges for patient diagnosis and prognosis. The development of next-generation sequencing and its decreasing costs promises to alleviate such issues by supplying personal genomic information at a moderate price. Here, we used crowdfunding as an alternative funding source to sequence the genome of Lil BUB, a celebrity cat affected by rare disease phenotypes characterized by supernumerary digits, osteopetrosis and dwarfism, all phenotypic traits that also occur in human patients. We discovered that Lil BUB is affected by two distinct mutations: a heterozygous mutation in the limb enhancer of the *Sonic hedgehog* gene, previously associated with polydactyly in Hemingway cats; and a novel homozygous frameshift deletion affecting the *TNFRSF11A* (*RANK*) gene, which has been linked to osteopetrosis in humans. We communicated the progress of this project to a large online audience, detailing the ‘inner workings’ of personalized whole genome sequencing with the aim of improving genetic literacy. Our results highlight the importance of genomic analysis in the identification of disease-causing mutations and support crowdfunding as a means to fund low-budget projects and as a platform for scientific communication.

## Introduction

Rare diseases represent a challenge for the research community and a public health problem. Although each disease individually affects a small number of people (no more than 1 person per 1,000-2,000, depending on the country), more than 7,000 different rare diseases have been reported. Therefore, it is estimated that the collective number of cases can be as high as 30 million in Europe and 25 million in the USA, representing 5-10% of the population^1^. Research on individual rare diseases has been traditionally neglected in favor of other, more common, medical conditions. Consequently, patients affected by these conditions often receive inadequate diagnostic and medical support, leading to so-called “diagnostic Odysseys”.

Comprehensive genetic testing holds promise to help such patients by identifying the cause of their disease, as well as enabling more informed, personalized treatments and prognoses. However, despite more than 10 years since the sequence of the human genome was revealed, our capability to interpret the effects of genomic variation remains limited. As a result, testing of candidate loci identifies disease-causing mutations for only a fraction of patients, and leaves a substantial number of unsolved cases. The increasing efficiency and decreasing cost of genome sequencing are expected to increase the number of available genomes, promising to improve the identification of disease-causing variants^2^ and further enhance our knowledge of the human genome. However, the next frontier lies on the inevitable challenges of evaluating hitherto unknown variants^3^, as well as communicating the potential and challenges of sequencing efforts to the patients and the general public, who often have no formal training in genetics, potentially leading to uncertainty and misunderstandings in their interpretation of the results^4–6^.

Animals provide a valuable resource in this framework to better understand genetic diseases, with domesticated animals and pets in particular displaying a rich tapestry of traits that often share the same genetic cause as in human^7–9^. One such example is Lil BUB, a pet cat with several congenital malformations (**Figure 1**), whose special appearance and social media presence has granted her international fame and a fandom across the globe. Lil BUB was diagnosed with infantile malignant osteopetrosis, a very rare condition that, in humans, affects 1 in 250,000-300,000 births. The disorder is caused by an impairment of osteoclasts, leading to an imbalance between the processes of bone resorption, performed by such cells, and bone formation, controlled by osteoblasts^10,11^. Together, these two cell types constantly remodel bone tissue throughout life to ensure its proper growth, mechanical stability and physiological functions. Thus, impairment of osteoclast differentiation or loss of resorptive activity can lead to the accumulation of bone tissue, causing increased bone density, susceptibility to fractures, as well as several traits that are secondary to those bone changes. Genetically, the disease is heterogeneous and causal mutations for the recessive, more severe form has been found in seven genes, which differ somewhat in their clinical manifestations and patient prognosis^10^. Currently, the only curative therapy is transplantation of hematopoietic stem cells, although alternative treatments such as gene therapy or cytokine-replacement therapy have been reported^10^. In cats, reports of osteopetrosis are rare^12,13^, and no examples of the congenital form are known, although it has been shown that osteopetrosis can be acquired through infection with retroviruses^14,15^.

**Figure 1.**
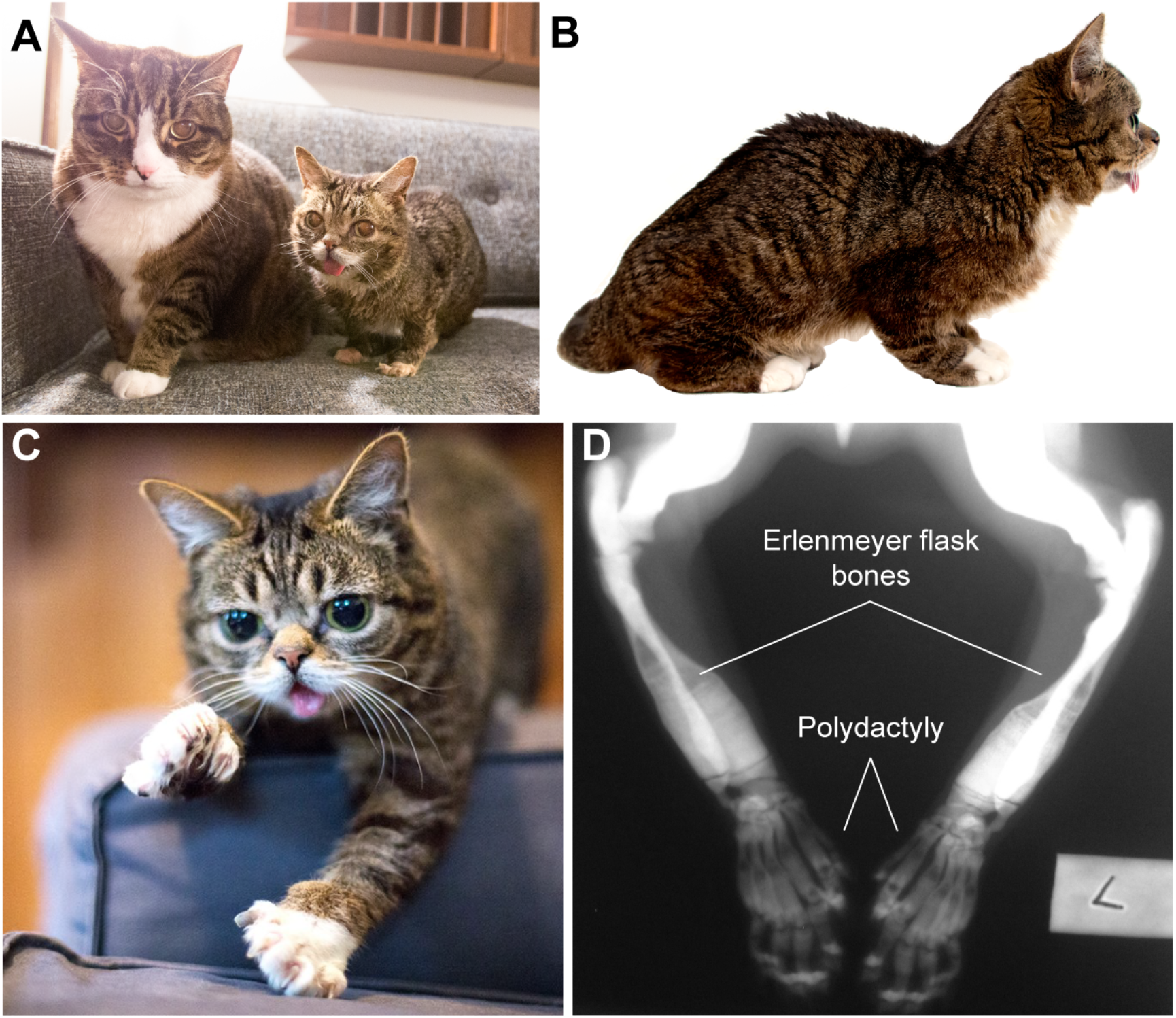
Phenotypic traits of Lil BUB. (A) Appearance of Lil BUB shows small size compared to an average adult domestic short haired cat (B) Short skull and body size, as well as protruding tongue shown in lateral view (C) Polydactyly of the front paws (D) Radiographs of forelimbs show high bone mineral density and Erlenmeyer flask shape of long bones as well as extra digits

In addition, Lil BUB presents preaxial polydactyly, a condition characterized by the presence of an extra digit on the thumb side of the paw. This trait is not uncommon in cats, with a high occurrence in several breeds (*e.g.* Maine Coon, Pixie bob) and multiple outbred cats^16,17^. Three mutations on chromosome A2, in the regulatory (enhancer) region of the *Sonic hedgehog* gene, have been previously identified as dominant causal variants^18^. Of these, one is known to be prevalent in North American cats, including in a colony of polydactyl cats on Ernest Hemingway’s estate on Key West^19^. Additional mutations on this enhancer and the associated polydactyly have been reported to occur in chicken, dog, mouse or human^18,20,21^, highlighting the evolutionary conservation of this developmental pathway.

The combined manifestation of congenital osteopetrosis and polydactyly, as observed in Lil BUB, is quite remarkable and has never been reported previously. To identify the underlying cause of the observed phenotypes, we performed a comprehensive genetic analysis for Lil BUB, and found that she is affected by two distinct mutations: a heterozygous mutation on the limb enhancer of the *Sonic hedgehog* gene; and a frameshift deletion affecting the *TNFRSF11A* (*RANK*) gene. We considered this a test case for precision medicine, since it would allow a more precise diagnosis and prognosis for Lil BUB and a general improvement of our understanding of the disease’s etiology. Due to the small-scale characteristics of the project, we determined to use a crowdfunding approach to finance the research. Moreover, we decided to communicate the planning, execution, and analysis of this study to the broader public using a range on online media platforms, to raise awareness for rare diseases and explain the challenges, promises and steps involved in whole genome sequencing. Furthermore, our results highlight the importance of genomic analyses as a means to improve our understanding of developmental disease.

## Results

### Phenotypic description

Lil BUB was found as a feral kitten that was abandoned by her mother in rural Indiana, USA and was subsequently hand-reared. She was born with several malformations, mainly related to skeletal growth and development. At the age of 7 months, she stopped growing and remained the size of a kitten (adult weight: 1.8 kg; **Figure 1A-C**). This manifested as an extreme case of dwarfism, causing her limbs to be remarkably small in comparison to her body. From birth she was smaller than other cats and by 10 months her movements became severely restricted. Radiologic studies revealed opaque medullary cavities and widened metaphyses resulting in an “Erlenmeyer Flask Deformity” and bowing of long bones (**Figure 1D**). Multiple radiopaque and radiolucent transverse bands were present in the metaphysis, a typical sign of problems with bone resorption. Ribs and sternae were thickened and the epiphyses of the long bones were small, while adjacent physes were thin. Moreover, her skull and jaw remained underdeveloped, with her tongue and eyes displaying a normal size and thus protruding notably from the cranial cavities. The teeth did not erupt or are not present, except for one or two small crowns on the right maxilla. There is a complete absence of bony palate. Despite her jaws and teeth being underdeveloped, Lil BUB has no difficulty feeding and eats food (both wet and dry) with no problem.

Based on her features and the radiological observations, she was diagnosed with a severe form of osteopetrosis, displaying similar features as described in human patients. Although this condition often co-occurs with additional symptoms that affect hormone and blood homeostasis, thyroid screening showed normal T4 (3.8 µg/ml) and Ca^2+^ (9.5mg/dl) levels and bloodwork showed no abnormalities, although enlarged platelets were noted. Lil BUB displayed low alkaline phosphatase levels (<20 IU/L; normal range 23-107 IU/L), which indicates low bone formation. However, in patients affected with osteopetrosis, these levels are often normal or even elevated because of the increased bone surface and varying total osteoblast numbers, and are thus unreliable biomarkers for osteopetrosis.. Of note, Lil BUB received continuous treatment with an electromagnetic pulse device (Assisi loop) since the age of 22 months. Coinciding with the treatment, her condition improved, and Lil BUB regained her mobility.

In addition to the traits associated with osteopetrosis, Lil BUB displays preaxial polydactyly affecting both forelimbs and hindlimbs (**Figure 1C and D**). A complete duplication of a normal biphalangeal thumb is observed in all paws, resulting in 22 digits (6 on each forepaw and 5 on both hindpaws).

### Project funding and outreach aspect

Given Lil BUB’s unique combination of osteopetrosis traits and polydactyly, we aimed to investigate whether her condition was an unusual form of (feline) osteopetrosis that included polydactyly, or if she was simultaneously affected by two rare conditions. We opted to perform whole genome sequencing to address this question, because a candidate gene approach did not seem feasible. We estimated that the financial requirements needed for such an endeavor would be moderate, and therefore decided to finance the project via crowdfunding^22^, with an initial goal of $6,500. We reasoned that Lil BUB’s recurrent engagement with the internet public presented a compelling case for “citizen science” that would benefit from alternative funding approaches, while simultaneously providing a platform to communicate genomic analyses to the broader public.

For this purpose, we used the crowdfunding platform “Experiment.com”, a site that focuses exclusively on raising funds for research and science projects. During the fundraising period, our campaign attracted the attention of more than 25,250 people (**Figure 2A**). After the 45-day campaign, the project was successfully funded, raising a total of $8,225, 26% more than the initial goal. The project was supported by a total of 248 “backers”, representing a 1% of the page views at the crowdfunding site during this period. Backers donated $33.17 on average. During the campaign, page views and donations peaked when the project was mentioned by the official Lil BUB Facebook page or major media outlets, such as the Washington Post or Der Spiegel (**Figure 2A** and **Supplementary Table S1**).

**Figure 2.**
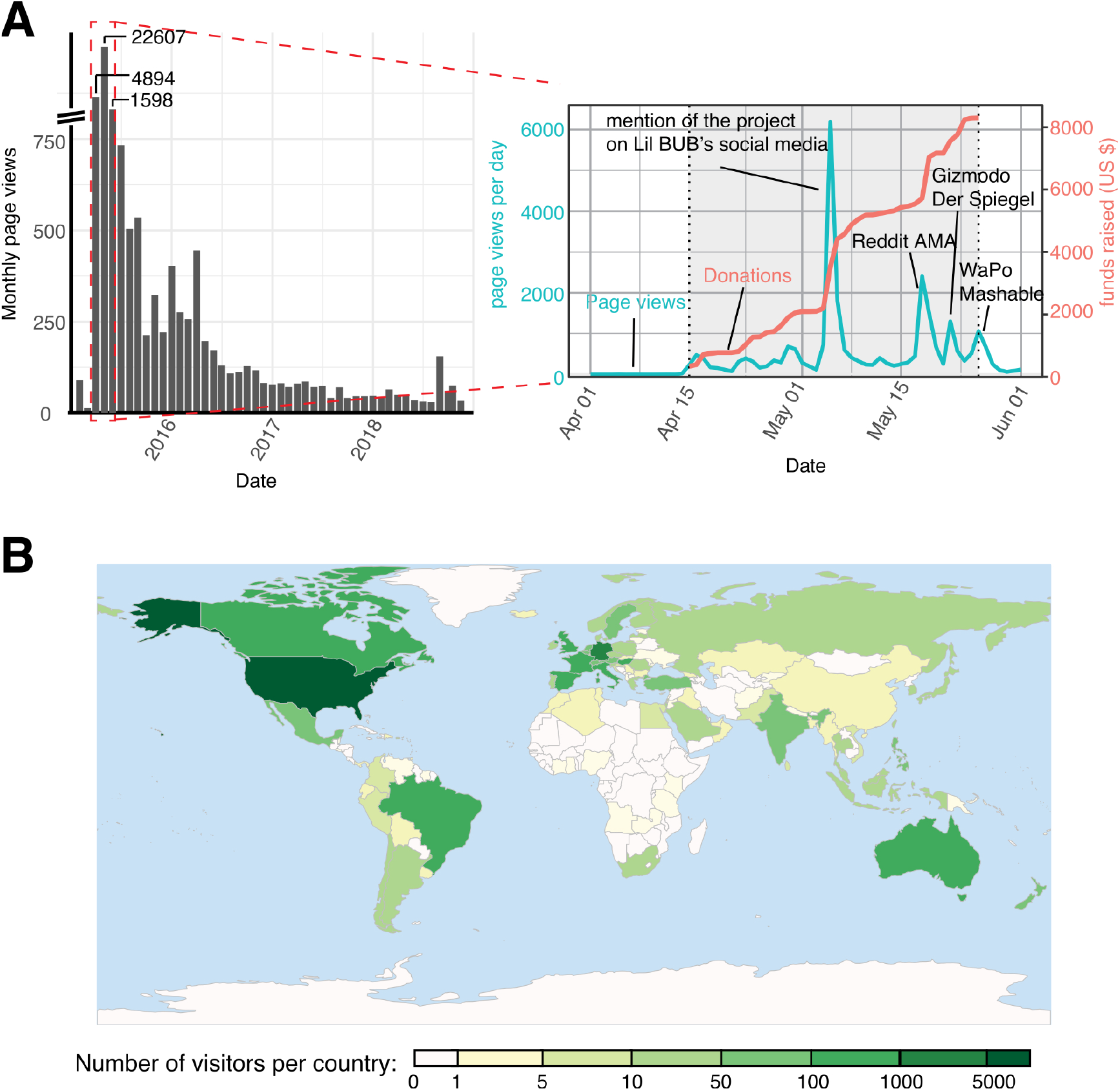
Web traffic on the homepages of the Lil BUBome. **(A)** Monthly number of visitors on the crowdfunding pages of the Lil BUBome at the experiment.com site between February 2015 and December 10, 2018 (left). It should be noted that for the three months with the highest number of visitors (April-June 2015) the height of the bars is not to scale. For these months we also show a daily breakdown of the site’s traffic (right), as well as the progress of fundraising during the crowdfunding period (April 15 to May 25, time highlighted in grey). On the days where we observed a spike in visitors, we also indicate what major media outlets or social media sites reported about the Lil BUBome. (B) Geographic distribution of visitors to the Lil BUBome’s Wordpress blog.

Beyond crowdfunding, we further used our social media networks to engage with the public and communicate the step-by-step progress of our research project. Our aim was to provide comprehensive explanations to the biology behind the project to a non-scientific audience, which represents most of the funders. One example was a Reddit Ask Me Anything (AMA) session that, over the course of 24 hours, allowed us to provide answers to specific project-related questions from the audience. We also produced YouTube video blogs to disseminate our results in a visual, more accessible format. As a consequence of these efforts, traffic on our sites continued after the crowdfunding period, with almost 11,000 additional visitors to our crowdfunding site, more than 5,000 views of our research updates and 15,000 visitors of our blog as of December 2018. We reached audiences from 109 countries on six continents, with approximately 50% of our websites’ traffic from the US and 11 other countries (US, Germany, UK, France, Canada, Hungary, Australia, Spain, Brazil, Italy and India) making up for 90% of our readers (**Figure 2B**).

### Genome sequencing and initial analysis of informative variants

To identify Lil BUB’s disease-causing mutations, we sequenced her genomic DNA using short-read sequencing technology. We mapped reads against the domestic cat reference genome (Felis_catus 6.2), obtaining ∼50X coverage and, through comparison with the reference genome, identified a total of ∼6 million SNVs and small insertions/deletions, as well as 424 larger structural variants (**Figure 3A-C**). However, the relatively large number of structural variants and their genomic distribution (**Figure 3B**) suggests that the great majority are likely false-positives due to problems in the genome annotation. We therefore did not follow up on those identified variants. Of note, we observed an excess of homozygosity throughout the entire genome, likely indicating inbreeding events over the course of multiple generations (**Figure 3D**).

**Figure 3.**
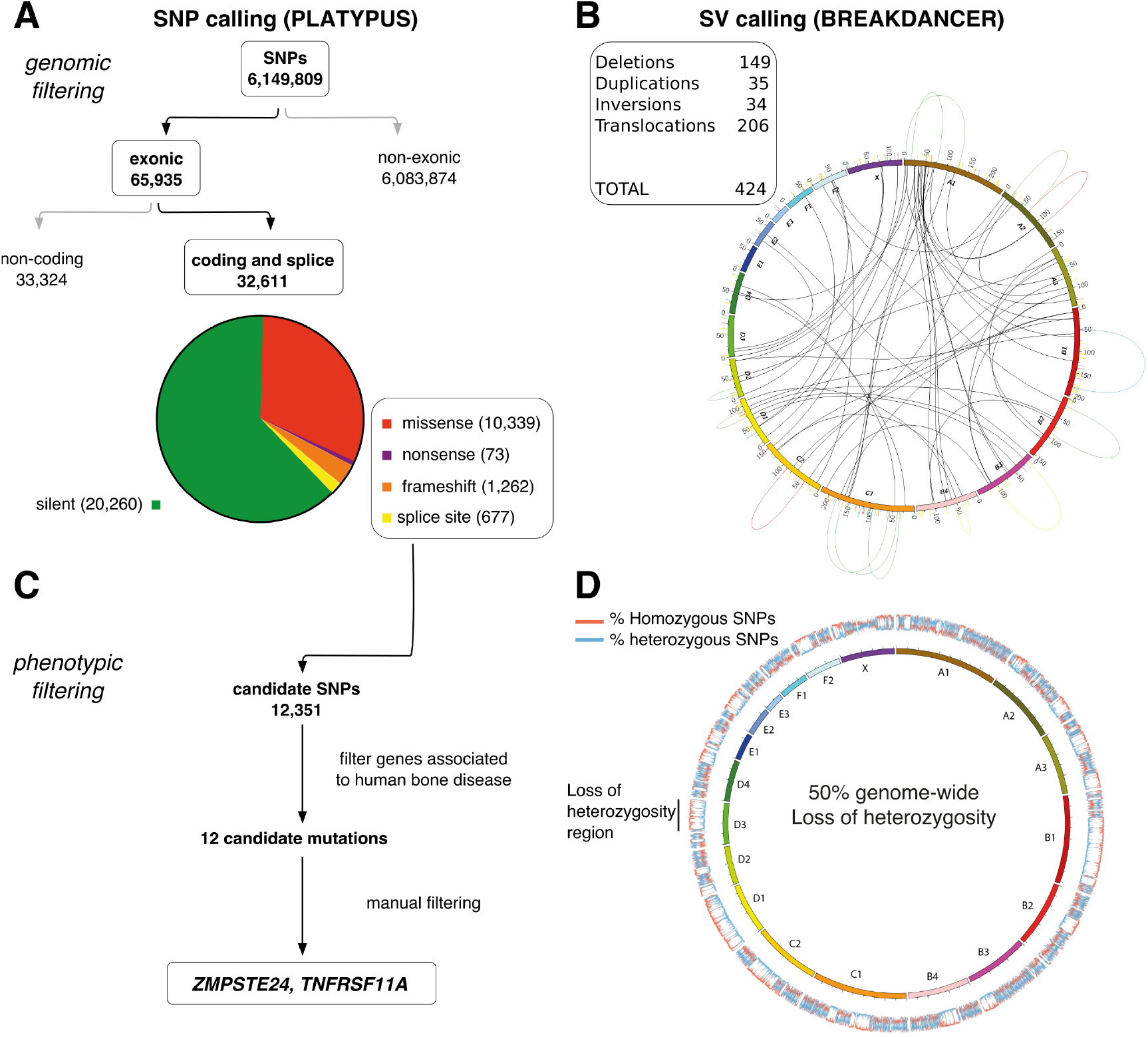
Variant detection in Lil BUB’s genome sequence. **(A, B)** Number and type of single nucleotide variants (A) and structural variants (B) detected in BUB’s genome relative to the reference cat genome, *Felis catus* v6.2, as determined using Platypus and Breakdancer tools, respectively. For each category, we indicate the number of observations in that category. The Circos plot for the structural variants shows the position and links of the detected structural variants. **(C)** Filtering of single nucleotide variants to identify candidate mutations for osteopetrosis. **(D)** Percentage of heterozygous and homozygous single nucleotide variants in 50kb windows along each chromosome show the genome-wide pattern of loss of heterozygosity.

As a first step of analyzing Lil BUB’s genome sequence, we looked at known variants previously linked to distinct phenotypical traits in cats (**Table 1**). In most cases, the studied variants matched with Lil BUB’s phenotypical characteristics. For example, Lil BUB does not harbor any of the mutations known to cause retinal atrophy, feline hereditary myopathy, hypertrophic cardiomyopathy or albinism. We found, however, that Lil BUB is a carrier for the variant c.123delCA in the *ASIP* gene that has been linked with black coat colour^23^. In addition, she carries two variants that affect the pattern of her coat’s tabby markings^24^. Such markings consist of light background hair, interspersed by darker hair in specific patterns, of which mackerel and blotched are the most frequent. In mackerel cats, the light and dark hairs form regular stripes. In contrast, the blotched pattern is characterized by disorganized stripes with the dark hair forming whorls. Lil BUB’s coat clearly displays the mackerel tabby pattern, although she is heterozygous for two distinct mutations (p.Trp841* and p.Thr139Asn) in the *Taqpep* gene that are associated with the blotched pattern. While our short-read sequencing cannot discern whether both variants are on the same chromosome, the co-occurrence of the two mutations with a mackerel pattern is consistent with published data, which found that 7 of 10 cats with the p.Trp841*/p.Thr139Asn mutations display this pattern^24^.

**Table 1.**
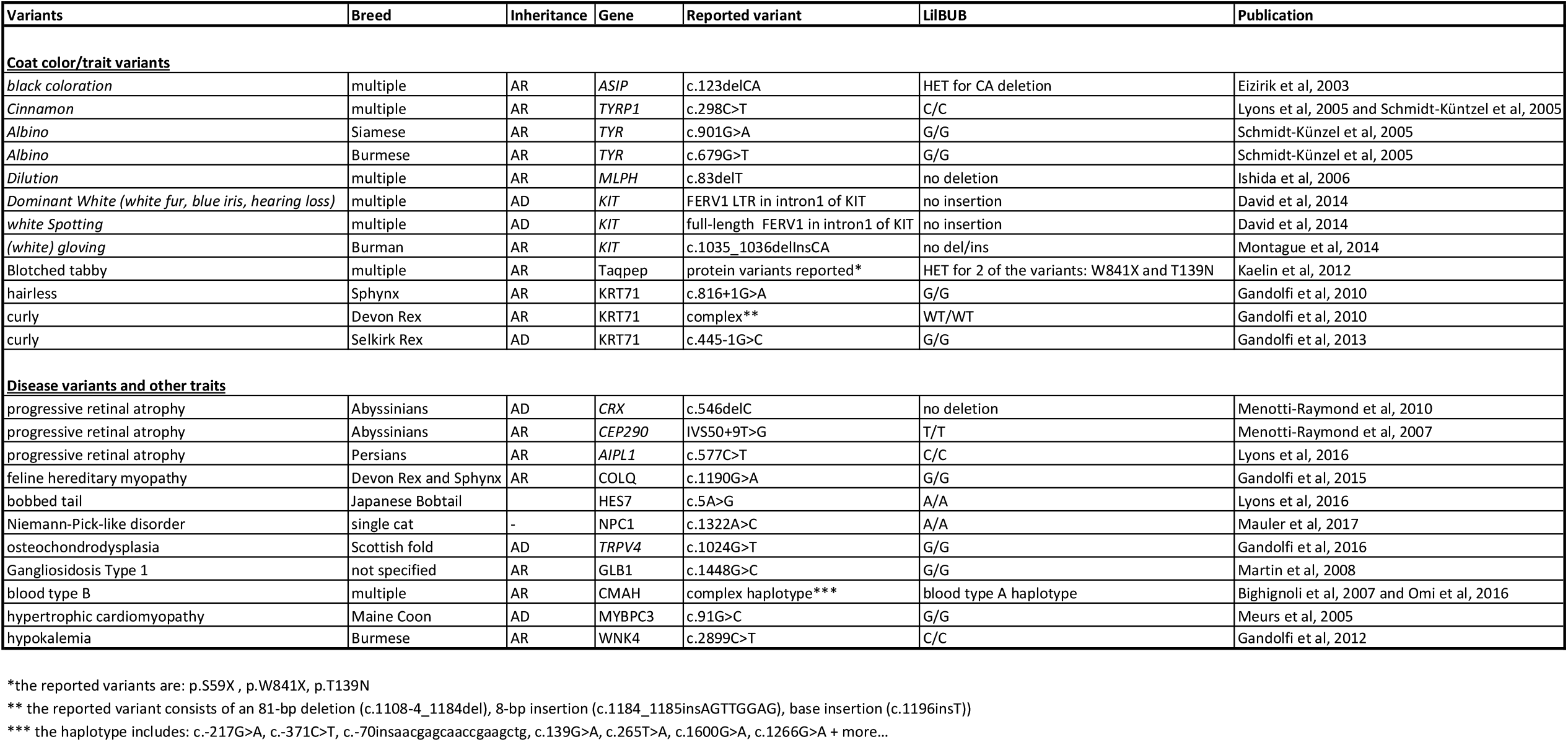
Status of known feline phenotype-associated variants in Lil BUB’s genomic sequence.

In addition to visible traits, her genetic findings were also consistent with molecular/physiological traits. For example, Lil BUB did not have the mutation associated with hypokalemia (intermittent reduced potassium levels) in Burmese cats. Consistent with this, she did not show any hypokalemic symptoms and her bloodwork displayed normal potassium levels (4.3 mmol/l; clinical signs appear below 3 mm/l)^25^. Similarly, we found Lil BUB to have the haplotype associated with blood type A in cats^26,27^. While her blood type was unknown at the time of our initial sequencing analysis, further tests confirmed that she indeed has blood type A. A major unexpected finding from our initial variant analysis was that, despite her white gloving phenotype (white fur only on her paws), she did not carry the variant known to cause gloving in the Birman breed. Such result suggests the existence of additional uncharacterized variants for gloving, an hypothesis also supported by observations in other breeds, such as the Ragdoll, displaying “mitted” patterns in the absence of the Birman mutation^28^.

### Identification of disease-causing mutations

Having performed this initial genotype-to-phenotype characterization, we next sought to identify the mutation(s) responsible for Lil BUB’s osteopetrosis and polydactyly traits. Initially, we chose a targeted approach and specifically analyzed the ZRS enhancer of the *Sonic hedgehog* gene, a genomic region associated to the appearance of polydactyly across mammals. We found that Lil BUB was heterozygous for an A to G mutation at chrA2:167,313,488, a variant previously described to cause the appearance of additional digits in Hemingway cats. We verified both the nature and the heterozygous state of the mutation using Sanger sequencing (**Figure 4A and B**). The affected enhancer is well-known for its association with polydactyly, with mutations in humans, mouse and cats identified in different parts of the element, which cause ectopic expression of *Sonic hedgehog* during embryonic limb development, subsequently leading to the formation of additional digits (**Figure 4C and D**).

**Figure 4.**
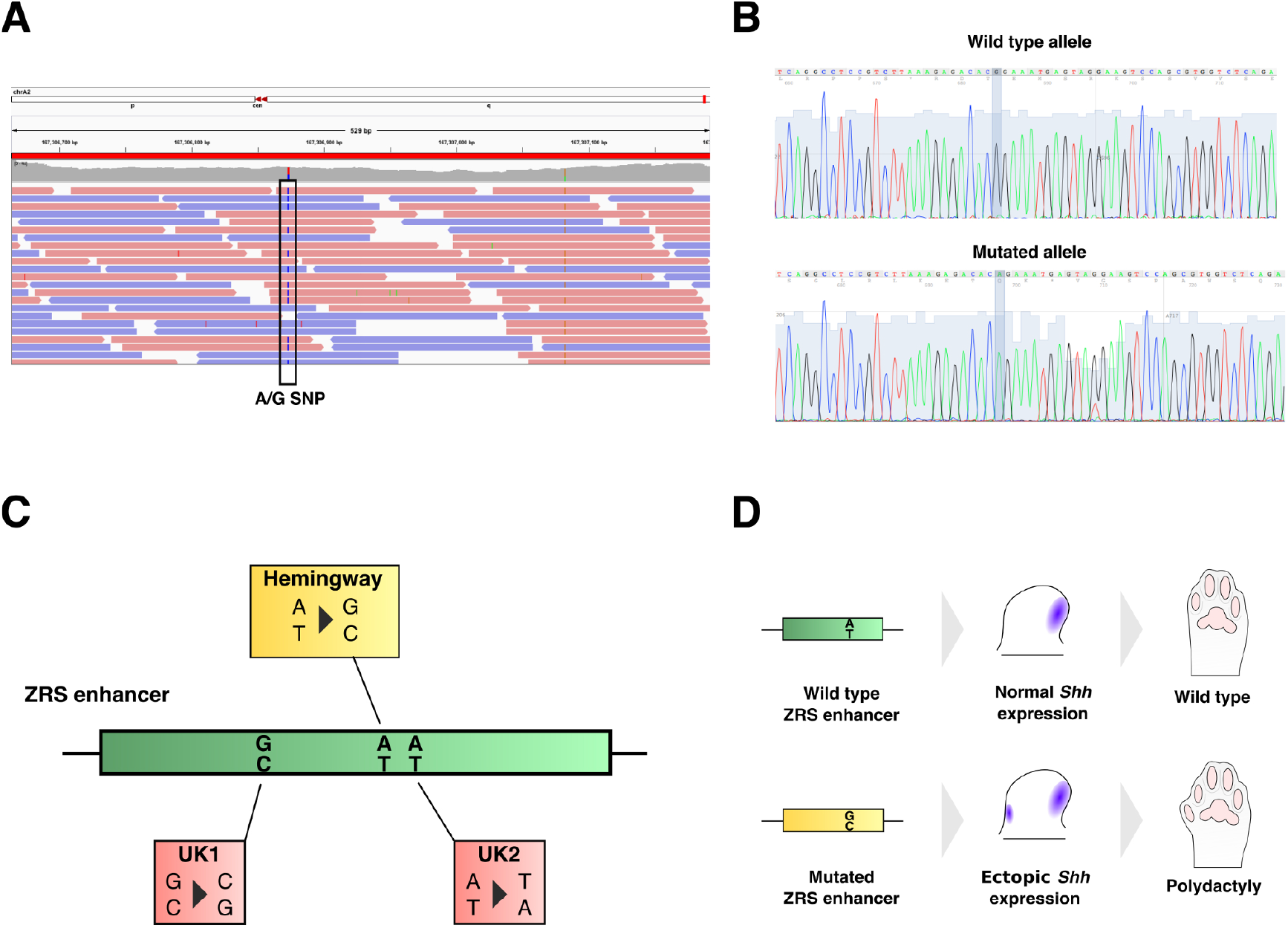
Polydactyly-associated mutations in the *Sonic hedgehog* limb enhancer. **(A)** A heterozygous A to G mutation at chrA2:167,313,488, in the limb enhancer of the *Sonic hedgehog* gene, as identified by whole genome sequencing in Lil BUB’s genome sequence. **(B)** Verification of Lil BUB’s mutation by Sanger sequencing, which showed the presence of a wild type allele (top), as well as a mutated allele (bottom). **(C)** Distribution of known polydactyly-associated mutations in the *Sonic hedgehog* limb enhancer in cats. **(D)** Developmental mechanism for polydactyly: The Hemingway-variant of the limb enhancer causes ectopic expression in the anterior part of the developing limb bud, giving rise to additional digits.

Having identified a likely candidate for the polydactyly phenotype, we next searched for causes underlying her osteopetrosis. Although we initially considered the possibility that retroviral infection may have given rise to Lil BUB’s osteopetrosis, she tested negative for both feline leukemia virus (FELV) and feline immunodeficiency virus (FIV). In addition, several healthy siblings were born in the same litter, thus eliminating this as a plausible causal mechanism. We therefore focused on the analysis of her genome sequence. We filtered Lil BUB’s variants in order to retain SNVs and inDels that affect genes (∼32k variants) and specifically focused on those with a known role in bone mass regulation and/or osteoclast differentiation in humans (**Supplementary Table S2**). Having thus narrowed the list to 12 candidate variants (**Table 2**), we further eliminated seven of these based on read coverage, mapping quality, and sequence complexity. We discarded three additional exonic variants, as manual inspection suggested that they were in fact intronic. This was subsequently confirmed by gene annotations in the genome builds 8.0 and 9.0.

**Table 2.**
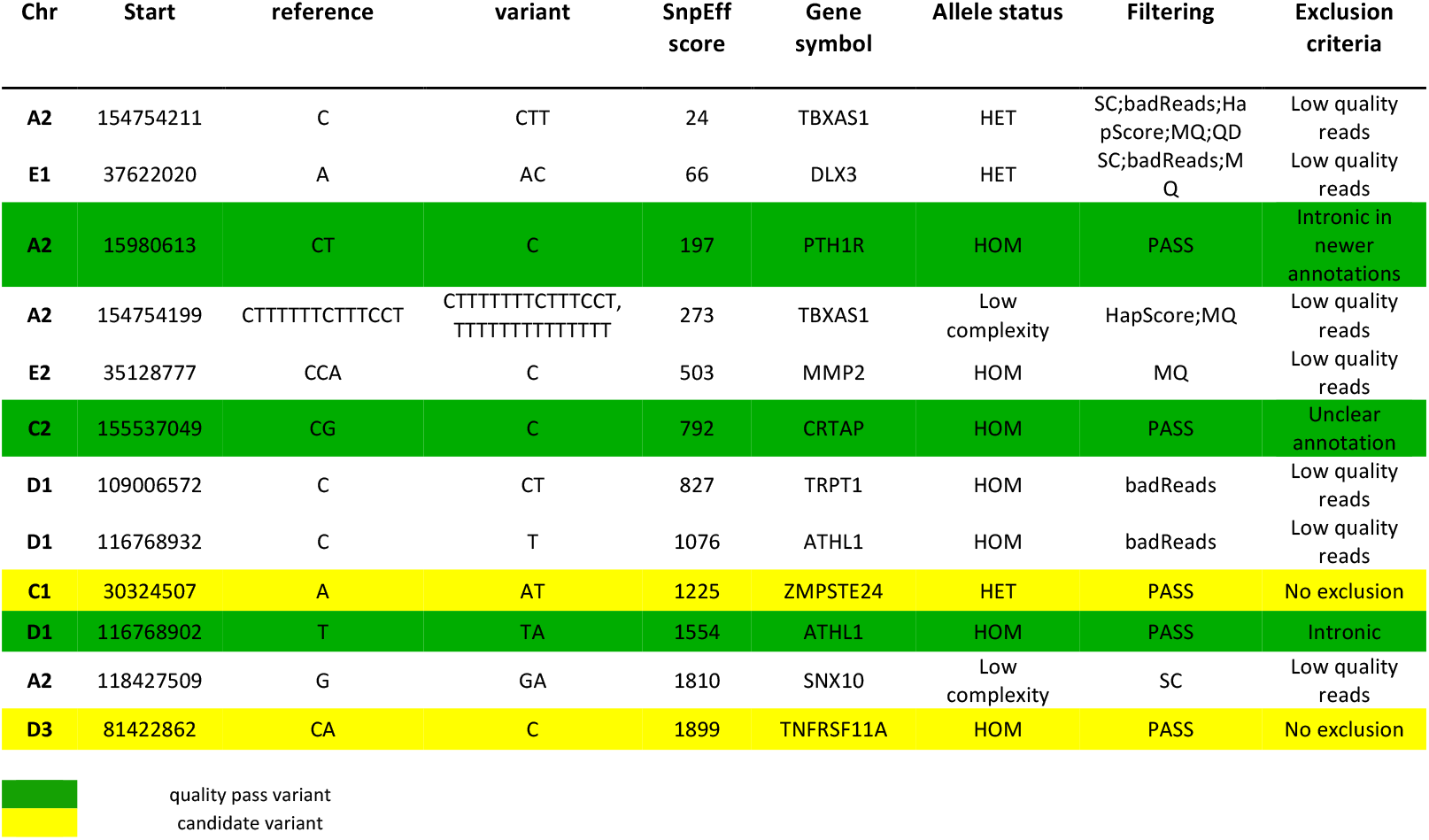
Shortlisted candidate genes for Lil BUB’s osteopetrosis. Yellow indicate variants that pass quality filters but are discarded due to unclear annotation in cat genome. Green indicate variants that pass quality filters and further retained for examination.

Of the two remaining variants, one involved *ZMPSTE24*, a gene that can cause mandibular defects in both human and mice when mutated^29–31^. While Lil BUB’s jaw is underdeveloped, providing superficial resemblance to the human and mouse *ZMPSTE24* loss-of-function phenotypes, diverse evidences suggest that it is not the major causal mutation for her traits. First, there are multiple features, typically associated with human and mouse *ZMPSTE24*-deficiency, that are inconsistent with Lil BUB’s phenotype. These include spontaneous bone fractures due to decreased bone density, lipodystrophy (general fat loss, although this feature might be secondary to feeding difficulties), atrophy of the skin and loss of hair or sparse, brittle hair^29,30,32,33^. Second, the variant identified in Lil BUB (a T insertion at chrC1:30,324,507, causing a frameshift in the protein) appears to be in a heterozygous state. This contrasts with other studies showing that affected human patients are homozygous or compound heterozygous, while heterozygous individuals are reported to be healthy^29,33,34^. In addition, heterozygous *Zmpste24* knockout mice are either indistinguishable from their wild-type littermates^32^ or develop much milder phenotypes^30^. For these reasons, we consider unlikely that Lil BUB’s *ZMPSTE24* mutation is causative for her skeletal malformations.

In contrast, several lines of evidence support that the mutation affecting the gene *TNFRSF11A* is likely the disease-causing variant for Lil BUB’s osteopetrosis. *TNFRSF11A* codes for a transmembrane receptor, RANK, that is expressed on osteoclasts and osteoclast precursors. Activation of this receptor by its ligand RANKL is necessary for osteoclast differentiation, survival, and function, and is therefore essential for the process of bone resorption. The identified mutation is a homozygous deletion of an A in a CAT sequence in exon 8 (c.806delA; **Figure 5A**), causing a frameshift and subsequent premature truncation of the protein (p.His272Leufs*16; **Figure 5B**). This truncation removes the majority of RANK’s intracellular domain, including two regions that are essential for oligomerization of the receptor and signal activation, thus rendering the protein dysfunctional^35^. Due to the central role of RANK in osteoclast function, variants that truncate murine or human RANK cause autosomal-recessive osteopetrosis, characterized by a complete absence of osteoclasts (**Figure 5C**). Affected individuals display early onset osteopetrosis, sharing several phenotypic features with Lil BUB, such as growth arrest and small stature, Erlenmeyer flask bones, exophtalamus (protruding eyes), dysmorphologies of the skull and jaw, and – in mice – failure of tooth eruption^36–40^. Two variants in particular, the human p.Gly280* and p.Trp434* mutations, are of interest as they also truncate the intracellular domain of RANK and biochemical assays of the deleted regions confirmed their involvement in RANK function^41,42^. Moreover, the identified variant was not present in 131 cats sequenced by the 99 Lives Consortium – none of which has osteopetrosis (data accessed August 2018). Taken together, these data strongly suggest that the homozygous, truncating *TNFRSF11A/*RANK variant is the causative mutation for the osteopetrotic phenotype observed in Lil BUB.

**Figure 5.**
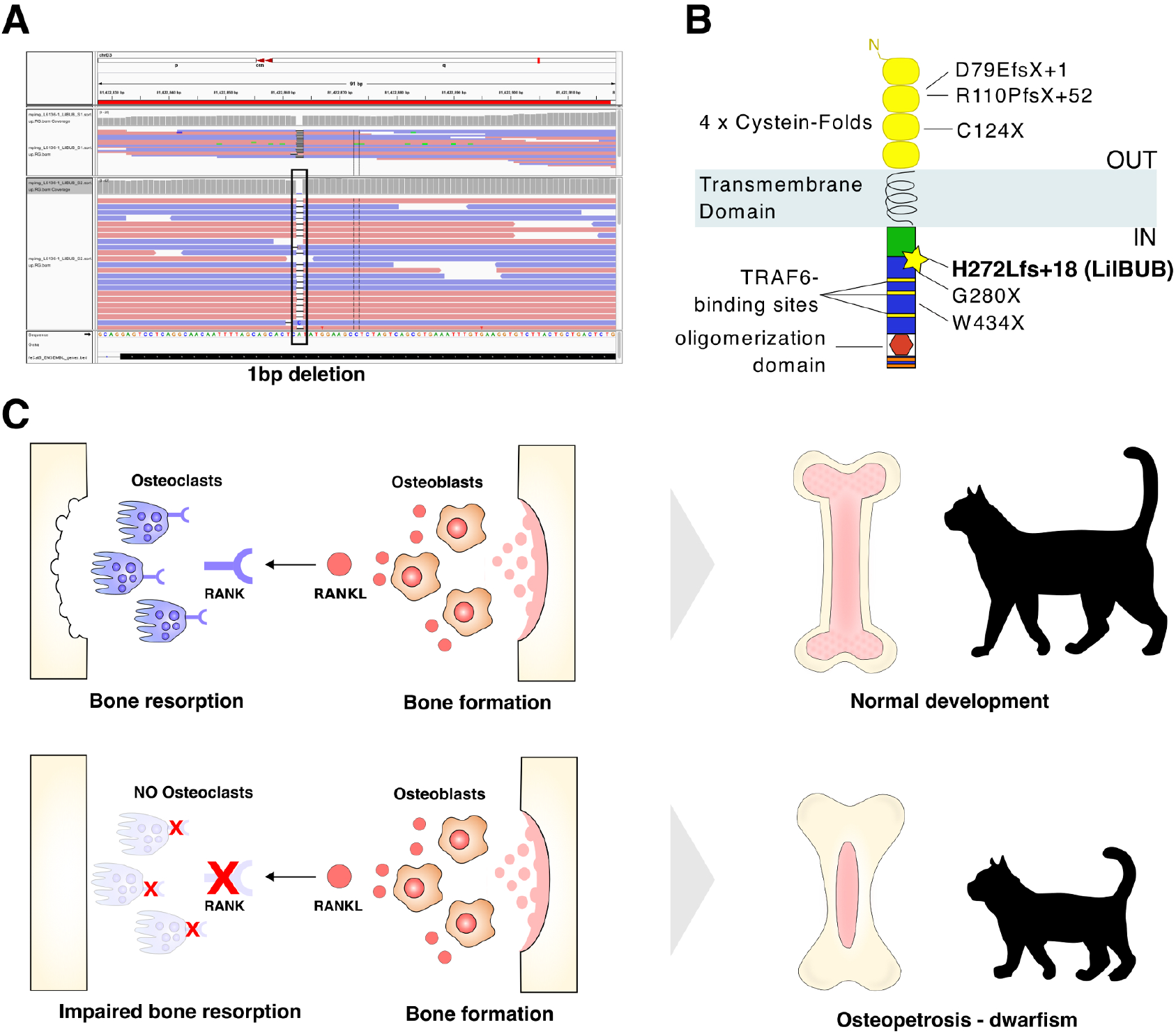
Osteopetrosis-associated variants in *TNFRSF11A*/RANK. **(A)** A homozygous 1-basepair deletion at chrD3:81,422,853, in exon 8 of *TNFRSF11A* (which codes for the protein RANK), as identified by whole genome sequencing in Lil BUB’s genome sequence. **(B)** Schematic structure of the RANK protein and known human and mouse mutations. The position of Lil BUB’s mutation is indicated by the star, and the two functional domains that are deleted due to the early truncation of the protein (the STAT-6 binding domain and the oligomerization domain) are also shown. **(C)** Developmental mechanism for osteopetrosis: bone formation and bone resorption are maintained by a balance of osteoblasts and osteoclasts, which express RANK-ligand (RANKL) and its receptor (RANK), respectively. A mutation that abolishes the function of RANK causes impaired bone resorption and thereby osteopetrosis and dwarfism.

## Discussion

Genetic studies in livestock and pet animals can provide valuable insight into human disease, due to the evolutionary conservation of mammalian body plans and their gene regulatory networks^43,44^. Domesticated animals, in particular, often display a vast array of readily observable phenotypes as a result of inbreeding^45,46^, which causes a gradual increase in homozygosity and the accumulation of recessive traits. Additionally, rare spontaneous mutations that mimic human conditions have also been documented^20,47,48^. Recent improvements in sequencing technologies have boosted our capacity to analyze entire genomes, expanding the potential to identify pathogenic mutations and constituting an important step towards disease management. Here, we demonstrate how personalized genomics and the conservation of mammalian disease phenotypes can be effectively linked, by sequencing the genome of a pet cat, Lil BUB, and identifying likely causal mutations of her polydactyly and osteopetrosis.

Lil BUB displays an unusual combination of polydactylous and osteopetrotic traits. However, mapping the genetic mutation(s) responsible for these traits through breeding or pedigree analysis was not possible, as she is spayed and her parents unknown. Therefore, we opted to perform whole genome sequencing to identify potential causal variants. By doing so, we identified approximately 6 million single nucleotide variants in Lil BUB compared to the reference genome, obtained from a female Abyssinian cat. This number of variants is similar to that observed in other breeds sequenced at high coverage^28^. Interestingly, we observed a substantially higher degree of homozygosity throughout Lil BUB’s genome than expected, based on prior studies in both inbred and random populations. In such studies, average heterozygosity measurements at the population level ranged between 0.51 and 0.65 (based on microsatellites) or 0.53-0.85 (based on short tandem repeat analysis^49,50^). Hence, our data suggests that multiple rounds of inbreeding occurred in Lil BUB’s ancestry, either as a result of a small cat population at her place of birth or because her ancestry is partially of a certain cat breed.

In the case of the osteopetrosis, we excluded the possibility that retroviral infection may have given rise to the disease, and identified a mutation in *TNFRSF11A*, a gene well known to cause such abnormalities. The identified mutation was homozygous, consistent with the autosomal recessive osteopetrosis subtype typically associated with *TNFRSF11A*/RANK (**Figure 5**). Beyond the phenotypical similarities between Lil BUB and mice or humans with RANK mutations, the identified variant was absent in all other tested cats of the 99 Lives Consortium, further supporting a causal relationship. Unfortunately, functional validations, such as an *in vitro* osteoclast differentiation assay, are not possible in this case, given the large blood sample requirements.

However, despite the overall similarity in phenotype with human and mouse *RANK*-associated osteopetrosis, we noted some differences: while the spleen is typically enlarged in humans and mice, no hepatosplenomegaly was observed in Lil BUB’s case. Furthermore, human patients affected by *RANK* mutations often display severe visual impairment, although Lil BUB’s vision seems to be unaffected. It is possible, however, that the described visual impairment is human-specific, as it has not been reported in mice^38–40^. Other Lil BUB’s features, such as severely hypoplastic jaw bones, have not been reported in human patients with RANKL-related autosomal recessive osteopetrosis, but pronounced mandibular hypoplasia was observed in cattle with *CLCN7*-related osteopetrosis, where it was paralleled by gingival hamartomas^51^. Thus, skull bones seem to be especially prone to species differences in autosomal recessive osteopetrosis. In addition, *RANK*-associated osteopetrosis can be linked with hypogammaglobulinemia and an increased risk of infections in some patients^36,37^. Although such parameters were not measured in Lil BUB’s bloodwork, the absence of remarkable recurrent infectious processes during her life suggests normal immunoglobulin levels. Lil BUB’s low alkaline phosphatase levels are notable since the enlarged osteopetrotic bone surface should lead to higher numbers of osteoblasts producing this enzyme. RANK has recently been identified as a coupling factor, which can stimulate osteoblast activity through binding to RANKL, thereby initiating reverse signaling^52^. Thus, a loss of RANK protein might explain this unusually weak bone formation, although cannot discern whether the identified frameshift mutation leads to a truncated, partly functional RANK protein or to a complete loss.ç

Currently, the only known therapy for autosomal recessive osteopetrosis in humans is bone marrow transplantation, which has not been required in the case of Lil BUB due to the favorable evolution of her condition. Such improvement was noticed soon after receiving regular sessions of pulsed electromagnetic field therapy. However, currently a scientific basis for the use of this method to treat the disease is missing and, to date, no other validated cases of feline osteopetrosis exist to verify if the observed effects are reproducible.

In contrast to osteopetrosis, polydactyly is a frequently observed trait in cats and the variant identified in Lil BUB has previously been documented in Hemingway cats, as well as other North American polydactyl breeds^53^. This non-coding mutation is assumed to result in polydactyly due to ectopic expression of the signaling molecule *Sonic hedgehog* in the anterior part of the developing limb buds (where it is normally not expressed). Interestingly, Hemingway cats are polydactylous mainly on the forelimbs^53^, while all four paws are affected in Lil BUB’s case. However, polydactyly has highly variable expressivity, even within a colony. At the same time, morphological variation of polydactyl cats is often reduced within breeding lines^16,17,53^, suggesting the existence of possible modifier genes. Based on our data we cannot distinguish whether there is a genetic component for the four-pawed polydactyly in Lil BUB or if her phenotype is simply part of a natural spectrum associated with the Hemingway mutation.

We also identified an additional mutation (in the *ZMPSTE24* gene) that could potentially have modified the outcome of her skeletal traits, raising the possibility that other exist. We also eliminated non-exonic variants, which can also affect gene regulation and/or splicing and need to be considered if no plausible coding mutation is detected. In addition, given that gene annotation of the cat genome is still a work-in-progress, it is possible that some variants currently annotated as non-coding will later be re-assigned, or *vice versa*, as we noticed during the analysis of Lil BUB’s genome (see **Methods**). This underlines the importance of thorough and comprehensive genome annotation for personalized genomics.

Finally, the efforts presented in this paper were also part of a public outreach project to demonstrate the use of personalized genomics: we raised funds for sequencing through a crowdfunding campaign and reported on various steps of sequencing and analysis on Twitter, through Facebook, blog posts and Reddit, as well as YouTube videos. Given the rise of direct-to-consumer genetic testing both for humans and, more recently, for pets^54,55^, we hope that our public engagement activities contributed to a broader understanding of how genetic information is obtained and what insights can (and cannot) be gained for a single individual case with unknown ancestry or family medical history.

## Methods

### Ethics statement

Blood samples from Lil BUB were collected by a licensed veterinarian, with the consent of her owner. According to the framework defined by the Directive 2010/63/UE of the European Parliament and of the Council, no animal experimentation, only non-experimental clinical veterinary practice (taking biopsies and imaging for disease diagnosis) was performed.

### Sample collection, DNA extraction and sequencing

A sample of EDTA anti-coagulated whole blood was collected during a routine veterinary visit and genomic DNA was isolated using standard procedures. The quality and integrity of the DNA was assessed using the A260/280 ratio and agarose gel electrophoresis. Libraries were then prepared using Truseq sample prep for Next Seq Version2: DNA was sheared to 300bp and 500bp with Covaris according to the protocol provided by Illumina, adaptors were ligated to the two libraries, and correctly sized fragments excised from gel. We performed paired end 2x 150bp sequencing of libraries on an Illumina NextSeq 500.

### Variant calling and analysis of variants

We obtained a total of 144,172,364 and 310,362,266 reads for the two libraries (∼500bp and ∼250bp length, respectively). Reads were mapped against Felis_catus_6.2 using Bowtie2 and we were able to map 98.38% of reads, corresponding to 40x coverage. Non-uniquely mapping reads were placed in a random fashion by Bowtie2. We did not apply repeat masking and all mapped reads were used. Sequence read de-duplication, indel re-assembly and subsequent SNV calling was performed with Platypus^56^. In total, we identified 6.19 million SNVs and InDels, 4.4 million (65.4%) were intergenic and 1.75 million (34.6%) were located in annotated genes. Variants were visualized using IGV browser^57^. We used variant predictor SnpEff^58^ to determine the potential pathogenic effects of candidate osteopetrosis genes. To exclude that the identified *TNFRSF11A* variant was not simply a low frequency variant in cats, we interrogated the 99 Lives Consortium data: in August 2018 we downloaded all variants identified in *TNFRSF11A/RANK* (this corresponded to the January 2017 variant analysis). The D3:81,422,862 variant was not present in 131 cats from this dataset.

### Evaluation of coding variants

Due to the incomplete annotation of the cat genome, five variants affecting transcripts of bone disease related genes had to be evaluated individually. For this the most likely correct transcription and coding unit was evaluated by integrating all available gene prediction algorithms, sequence conservation, and expressed sequences in the UCSC genome browser for releases felCat5 and felCat8.0 of the *Felis catus* genome. Annotation inconsistencies of the *TNFRSF11A* transcript and coding sequence lead to ambiguities in the nomenclature of the frameshift variant. The most appropriate nomenclature is ENSEMBL c.806delA (ENSFCAT00000032280.1) and p.H270LfsX16 (ENSFCAP00000022567.1) in felCat5 or Augustus c.815delA (g12385.t1) and p.H272LfsX16 (g12385.t1_prot) in felCat8.

### Sanger sequencing of the ZRS region

A fragment containing the ZRS region was PCR generated using primers that amplify a 2.8kb region surrounding the 800bp ZRS (fwd_primer: CCTTGAAGTGGAAATCTCTCCTG, rev_primer: ATCAATTGCGTGAAAACTGCAAGGG). The obtained PCR products were subcloned into a pTA-vector and four individual subclones were sequenced using six sequencing primers that together covered 2.7kb of the region: catZRSeq1: TTGATGGGGTTTTCCTCGAAC, catZRSeq2: TCAGCTTTATAGGCCTTCCCAG, catZRSeq3: CAAGACGCAAACCGCGGAG, catZRSeq4: GGGCGGATGCAGAGCTTG, catZRSeq5: TTAGTGAGATATGAGTCCATTTTCTGT, catZRSeq6: GAGCATAGCACACGGTCT.

### Data availability

Sequencing data was deposited in the Sequence Read Archive with accession number PRJNA512113. It is available using the following link: https://www.ncbi.nlm.nih.gov/sra/PRJNA512113

---

**Table S1. A sample of media outlets that reported on about the sequencing of Lil BUB’s genome around the world.**

**Table S2. Genes with known role in bone mass regulation and/or osteoclast differentiation in humans.**

## Acknowledgements

We thank all the generous donors who contributed to our crowdfunding campaign, and without whom this project would not have been possible. We are also grateful for the guidance we received from experiment.com staff, especially Cindy Wu and Denny Luan, who helped us navigate through the funding process and aided our communication endeavors. We would further like to express our gratitude to our mentors, Stefan Mundlos, Gaël Yvert and Arjun Raj, who supported our work on this project. We would also like to thank Sarah Malek and Karun Kiani for providing critical feedback on the manuscript. Funding for the 99 Lives Cat Genome Sequencing Initiative has been provided by the Winn Feline Foundation and the George Sydney and Phyllis Redman Miller Trust (MT–13–010), the National Geographic Society Education Foundation (2P-14), the University of Missouri Gilbreath-McLorn Endowment (LAL), the Cornell Feline Health Center (ARB, MGC, RJT), contributions from the 99 Lives Consortium participants, donations from Associazione Nazionale Felina Italiana (ANFI), Zoetis, Orivet Genetic Pet Care, Langford Veterinary Services, the World Cat Federation, and public donations. We also wish to thank Illumina for providing a complimentary NextSeq 500 and Library Prep kit.

## Author contributions

D.M.I, D.G.L. and O.S. conceived the project, M.B. and A.W. performed disease diagnosis. M.B., U.W., B.T. and A.W. provided reagents. D.M.I., D.G.L and N.M. performed experiments, H.K., D.M.I., D.G.L. and O.S. analyzed the data. The 99 Lives Consortium provided the variant genetic database. D.M.I., D.G.L. and O.S. wrote the manuscript with feedback from all other authors.

